# A lack of population structure characterizes the invasive *Lonicera japonica* in West Virginia and across eastern North America

**DOI:** 10.1101/2023.03.01.530604

**Authors:** Craig F. Barrett, Cameron W. Corbett, Hana L. Thixton-Nolan, Biology 320 Class

## Abstract

Invasive plant species cause massive ecosystem damage globally, yet represent powerful case studies in population genetics and rapid adaptation to new habitats. The availability of digitized herbarium collections data, and the ubiquity of invasive species across the landscape make them highly accessible for studies of invasion history and population dynamics associated with their introduction, establishment, spread, and ecological interactions. Here we focus on *Lonicera japonica*, one of the most damaging invasive vine species in North America. We leveraged digitized collections data and contemporary field collections to reconstruct the invasion history and characterize patterns of genomic variation in the eastern USA, using a straightforward method for generating nucleotide polymorphism data and a recently published, chromosome-level genome for the species. We found an overall lack of population structure among sites in northern West Virginia, USA, as well as across sites in the central and eastern USA. Heterozygosity and population differentiation were both low based on *Fst*, analysis of molecular variance, principal components analysis, and cluster-based analyses. We also found evidence of high inbreeding coefficients and significant linkage disequilibrium, in line with the ability of this otherwise outcrossing, perennial species to propagate vegetatively. Our findings corroborate earlier studies based on allozyme data, and suggest that intentional, human-assisted spread explains the lack of population structure, as this species was planted for erosion control and as an ornamental, escaping cultivation repeatedly across the USA. Finally, we discuss how plant invasion genomics can be incorporated into experiential undergraduate education as a way to integrate teaching and research.

## INTRODUCTION

Invasive species cause billions of dollars in damage to habitats in the US and around the globe (Simberloff 2010). Yet, they provide important case studies in ecosystem dynamics and rapid evolution to new environments (Lawson-Handley et al. 2011). Traditional understanding of genetic diversity within invasive species focused on single introductions, subsequent genetic bottlenecks, and hypothesized low genetic diversity in the invasive range compared to that in the native range (see Tsutsui et al. 2000; Lee 2002; Frankham 2005; Estoup et al. 2016). While this was the case for many invasive species, the application of molecular markers have consistently identified similar or even higher levels of genetic variation in invasive populations compared to those in the native range (e.g., Frankham 1997; Kolbe et al. 2004). More recently, many studies have identified multiple introductions over space and time in invasive species, including admixture among originally isolated allele pools, and “bridgehead” introductions, whereby invasions occur in successive stages across regions or continents (Dlugosch and Parker 2008; Keller and Taylor 2010; Van Boheemen et al. 2017; Vallejo-Marín et al. 2021). Thus, the picture emerging from molecular genetic studies of invasive species is more complex than “traditional” hypotheses of invasion, and represents the interplay between history, dispersal, breeding system, source and recipient habitats, and several other factors (Sakai et al. 2001; Sutherland et al. 2021).

Information from digitized collections databases provides a useful tool for reconstructing invasion routes and history, trait variation over space and time, and physical collections provide genomic resources for spatiotemporal analysis of variation (e.g. Gallinat et al. 2018; Barrett et al. 2022; Bieker et al. 2022; Heberling 2022). In parallel, advances in genomic sequencing (RAD-seq, GBS, low-coverage whole genome sequencing, sequence capture, multiplexed amplicon sequencing) provide increased power over previous methods (allozyme variation, organellar gene/spacer sequencing, microsatellites) for studies of genetic variation and population structure, with broader representation of the genome for detecting both neutral and adaptive variation (Chown et al. 2015; Hamelin and Roe 2020; North et al. 2021). In plant biology, many of these technological advances have focused on crops or threatened/endangered species; relatively fewer have focused on invasive species (Barrett 2015; Hohenloe et al. 2020; North et al. 2021).

*Lonicera japonica* Thunb. is one of the most aggressive, invasive vines in eastern North America, yet is surprisingly not well studied from a genetic perspective across its globally invasive range, including North America (Schierenbeck 2004). This species forms dense mats, climbs trees and shrubs, outcompetes native vines and understory species, and causes tree mortality (Leatherman 1955; Evans 1982; Hardt 1986; Dillenberg et al. 1993). Traditionally, this species has been planted as a means of erosion control and as an ornamental, from which it is hypothesized to have escaped cultivation repeatedly. *Lonicera japonica* is highly attractive to diverse pollinators, with large nectar rewards, and the seeds are dispersed locally by birds and mammals, and by humans over greater distances (Luken 1996). *Lonicera japonica* is an obligate outcrosser but also propagates vegetatively by rerooting from stems, forming clonal ramets in many places (Leatherman 1955). At least 12 cultivars are known, but “Hall’s honeysuckle” is believed to be the most common and prolific, and further hypothesized to be the major player in invasion across the USA (Schierenbeck 2004).

Studies based on allozyme electrophoresis revealed low levels of genetic diversity in the southeastern USA (Schierenbeck et al. 1995; Schierenbeck 2004), yet genome-scale data and analysis are yet to be applied to quantify patterns of diversity in this species. A chromosome-level genome was recently published (Pu et al. 2020), providing a powerful resource for population genomics. In addition, several economical, technologically straightforward methods have recently been published based on amplicon sequencing for the generation of single nucleotide polymorphisms (SNPs), allowing genome-scale assessments of population-level variation that were not possible previously (e.g. Suyama and Matsuki 2015; Sinn, Simon et al. 2021). Thus, the tools and resources for the study of invasion genomics are now at hand for numerous species, including *L. japonica*.

It remains a challenge to develop undergraduate-level courses that capture the entire scientific process. Previous educational models or programs have identified positive impacts on student learning such as students gaining deeper insight into the research process or overcoming research-associated challenges within a positive, supportive environment (Lopatto et al. 2020). Many of these programs were focused on formative outcomes rather than tangible, summative outcomes such as students presenting their findings in a public environment (e.g. poster symposia; Wei and Woodin 2017, Campbell and Nehm 2017, Haskel-Ittah et al. 2020). A similar course-based undergraduate research experience (CURE) was developed for the invasive grass *Bromus inermis*, but focused primarily on ecological questions, and the course was designed for first-year undergraduate students (Laungani et al. 2018). To our knowledge, there has been minimal work with undergraduate students utilizing plant invasion biology to address questions of well-established invasive species, and thus we saw an opportunity to integrate CURE principles within an emerging and understudied field of research.

Our objectives were three-fold: 1) Mapping the invasion history of *L. japonica* in the USA using digitized herbarium specimen information, 2) quantifying patterns of genetic diversity and population structure across the eastern USA using genomic data, and 3) using plant invasion genomics as an experiential learning system in undergraduate education. We sampled 166 individuals across 16 localities in eastern North America (with a focus on northern West Virginia), and employed a straightforward, amplicon-based protocol (MIG-seq, or Multiplexed inter-simple sequence repeat genotyping) to quantify genomic variation in *L. japonica*. Our analysis yielded >1,500 SNPs, and revealed an overall lack of population structure for this invasive species in the eastern USA, suggesting a highly admixed gene pool. Lastly, we discuss how invasive species can be used as an effective teaching tool for integrating undergraduate research experiences.

## MATERIALS & METHODS

### Reconstructing invasion history with herbarium records

We created an animation using database records from herbarium specimens collected over the past two centuries. Specimen information was accessed through the Global Biodiversity Information Center (GBIF; https://www.gbif.org/, last accessed 17 February, 2023) with the R package rgbif v.3.7.5 (Chamberlain et al. 2023), and the animation was created in R following Barrett et al. (2022). A static representation was also created across six time slices of 30 years each, from 1880-2020 (except for the latter slice, which was 20 years from 2000-2020). Code for the animation and static maps (and the figures themselves) can be accessed via GitHub (https://github.com/barrettlab/2021-Genomics-bootcamp/wiki/2022-Biol-320-Lonicera-japonica-invasion-history-animation)

### Sampling, DNA extraction, and MIG-seq

Whole green leaves were collected from 166 individuals at 16 localities in the eastern and midwestern US (Table 1). At each sampling site, leaf samples were collected at least 10 m apart to avoid collecting tissue from the same ramet. One individual from each locality was pressed as a voucher specimen and deposited at the West Virginia University Herbarium. A large number of individuals were collected at the West Virginia University Earl Core Arboretum and in the City of Morgantown, WV; smaller samples were collected at other localities locally in WV and more broadly across the eastern US (Table 1). An ethanol- sterilized marker cap was used to punch an equal area of tissue (1 cm diameter) from each leaf, avoiding the midvein. The CTAB DNA extraction procedure (Doyle and Doyle 1987) was used to isolate genomic DNAs, using a modified 96-well extraction protocol. Briefly, samples were stored in 2 ml screw-cap tubes, frozen in liquid nitrogen, and pulverized with 3 mm steel bearings. DNA concentrations were measured with a plate reader (broad-range assay, Tecan Group, Ltd., Zurich, Switzerland) and diluted to 20 ng/ul in TE buffer (pH 8.0). MIG-seq amplicons were produced following the procedure in Suyama and Matsuki (2015), but modified for dual indexing. PCR conditions were as follows: 98°C 5 min, followed by 30 cycles of 98°C (30s), 48°C (30s), and 72°C (90s), with a final extension at 72°C for 5 min. PCR products were then diluted 1:50 in sterile PCR water, and used in a second round of PCR to add dual-indexed barcodes (File S1, https://doi.org/10.5281/zenodo.7686073). Cycle conditions were as follows: 15 cycles of: 98°C (10s), 54°C (15s), and 72°C (1m). PCR products were then quantified via NanoDrop spectrophotometry (Thermo Fisher, Waltham, Massachusetts, USA) and pooled at equimolar ratios. A single, two-sided PCR cleanup/size selection was conducted with Quantabio SparQ beads (Beverly, Massachusetts, USA) at bead to sample ratios of 0.8x and 0.56x. The resulted size-selected library pool was quantified with an Agilent Bioanalyzer (Santa Clara, California, USA) and with quantitative PCR, and sequenced at the West Virginia University Genomics Core Facility on two runs of an Illumina MiSeq using v.3 chemistry for 2 x 300 bp reads.

**Table 1.** Sampling locality and basic diversity information. N = sample size, Fis = inbreeding coefficient, Ia and r-barD = metrics of linkage disequilibrium with associated p-values based on 999 permutations.

| Locality | Latitude | Longitude | N | *Fis* | *Ia* | *P-value* | *r-barD* | *P-value* |
| --- | --- | --- | --- | --- | --- | --- | --- | --- |
| Allegheny, PA | 40.53315 | -79.781593 | 6 | 0.663 | 1.015 | 0.76 | 0.010 | 0.67 |
| Arboretum,<br>Monongalia, WV | 39.644729 | -79.976635 | 54 | 0.772 | 3.333 | **0.01** | 0.009 | **0.01** |
| Cheat Lake,<br>Monongalia, WV | 39.6715 | -79.847774 | 14 | 0.71 | 1.192 | 0.91 | 0.006 | 0.84 |
| Deckers Creek,<br>Monongalia, WV | 39.628758 | -79.949796 | 10 | 0.716 | 0.982 | 0.69 | 0.007 | 0.61 |
| Durham, NC | 36.024027 | -78.924629 | 6 | 0.712 | 2.003 | 0.26 | 0.017 | 0.22 |
| Fayette, PA | 39.797474 | -79.794005 | 3 | 0.637 | 2.031 | 0.17 | 0.023 | 0.17 |
| Jackson, WV | 38.82314 | -81.719958 | 3 | 0.589 | 7.534 | **0.02** | 0.076 | **0.02** |
| **Lawrence, PA** | 41.02137 | -80.446194 | 6 | 0.687 | 1.597 | **0.04** | 0.012 | **0.03** |
| **LittleFalls,**<br>**Monongalia, WV** | 39.553667 | -80.015538 | 8 | 0.693 | 0.855 | 0.62 | 0.006 | 0.55 |
| **Life Sciences**<br>**Bldg.,**<br>**Monongalia, WV** | 39.63786 | -79.95392 | 5 | 0.611 | 0.798 | 0.1 | 0.007 | 0.08 |
| **Morgantown,**<br>**Monongalia, WV** | 39.645545 | -79.980235 | 20 | 0.736 | 5.659 | **0.01** | 0.018 | **0.01** |
| **Oktibbeha, MS** | 33.450835 | -88.918471 | 3 | 0.644 | -0.964 | 0.97 | -0.012 | 0.96 |
| **Preston, WV** | 39.65972 | -79.791026 | 7 | 0.663 | 3.832 | **0.02** | 0.019 | **0.01** |
| **Shelby, TN** | 35.143333 | -89.986183 | 5 | 0.662 | 1.056 | 0.05 | 0.011 | 0.04 |
| **Star City,**<br>**Monongalia, WV** | 39.683194 | -79.961472 | 8 | 0.689 | 1.618 | **0.04** | 0.012 | **0.04** |
| **Uffington,**<br>**Monongalia, WV** | 39.587624 | -80.002644 | 6 | 0.668 | 0.678 | 0.25 | 0.007 | 0.2 |

### Read processing, mapping, and SNP calling

Reads were processed using a dedicated pipeline designed for amplified ISSR fragments (https://github.com/btsinn/ISSRseq/wiki). Briefly, reads are trimmed using BBDUK (http://jgi.doe.gov/data-and-tools/bb-tools) and mapped to an indexed reference sequence (here, the *Lonicera japonica* genome, NCBI BioProject accession PRJNA794868; Pu et al. 2020). Resulting BAM alignment files were sorted and PCR duplicates were removed with SAMTOOLS v.1.7-13 and PICARD v.3.0.0, respectively (Danecek et al. 2021, http://broadinstitute.github.io/picard). BAM files were then analyzed with GATK4 v.4.2, specifically using GATK’s “Best Practices” filters, using HaplotypeCaller to realign around indels (Van der Auwera et al. 2013; Poplin et al. 2017; Van der Auwera and O’Connor 2020). The resulting variants, called across all samples, were output as a .vcf file. This file was further filtered on missing data (for sites and individuals) and minor allele frequencies (removing minor allele sites with frequency < 0.05), and further thinned to keep only a single SNP per locus (minimum distance = 1,000 bp) with TASSEL5 v.5.0 (Bradbury et al. 2007) and PLINK v1.90b6.24 (Purcell et al., 2007).

#### Population diversity and structure analyses

Population genetic analyses were conducted with SambaR v.1.08 (De Jong et al. 2021), ADEGENET v.2.1.0 (Jombart et al. 2008), HIERFSTAT v.0.5.11 (Goudet 2005), POPPR v.2.9.3 (Kamvar et al. 2014), and SNPRelate v.1.32.2 (Zheng et al. 2012), following Sinn, Simon et al. (2021). Inbreeding coefficients (*Fis*) were calculated in SambaR and population differentiation (*Fst*) metrics were calculated with POPPR. Analysis of Molecular Variance was performed with HIERFSTAT, testing the significance of the components of variation with 999 permutations. Principal Components Analysis was conducted with SNPRelate. Heterozygosity values (observed, *Ho*, and expected, *He*) for each locality were calculated with SambaR and POPPR. Discriminant Analysis of Principal Components (Jombart et al., 2010) was conducted in ADEGENET, choosing “k,” or the number of ancestral genomic population clusters, using the Bayesian Information Criterion to select among different k-values. The cross-validation method in ADEGENET was used to choose the optimal number of Principal Components in the analysis. A multilocus genotype network was also constructed with ADEGENET. Finally, a dendrogram based on Nei’s Genetic Distance was created, grouping by locality, with the ‘aboot’ function in POPPR, with 1,000 bootstrap pseudoreplicates. All plots were created with the ggplot2 v.3.4.1 (Wickham 2016), ggpubR v.0.6.0 (Kassambara et al. 2020), and pheatmap v.1.0.12 (https://github.com/raivokolde/pheatmap) packages for R.

## RESULTS

### Invasion history

Plotting of historical herbarium records over six time slices revealed a rapid colonization of *Lonicera japonica* across the eastern USA (Fig. 1; Animation: https://github.com/barrettlab/2021-Genomics-bootcamp/wiki/2022-Biol-320-Lonicera-japonica-invasion-history-animation). By 1880, this species was present around New York City, in upstate New York, and in northern Virginia and Maryland. By 1910 it had spread to Pennsylvania, the Carolinas, Florida, Georgia, Arkansas, Missouri, Texas, and as far west as California (a single record, near Lake Tahoe on the Nevada border). By 1940 the spread continued in the northeastern USA, southeastern USA, midwestern USA, and within California. By 1970 it had spread northward into New England and the northern Midwest, and also into southern California, Arizona, and New Mexico. By 2000 it had invaded nearly all of the eastern USA except Maine, most of California, and by 2020 it had spread into Colorado, Oregon, and Idaho.

**Figure 1.**
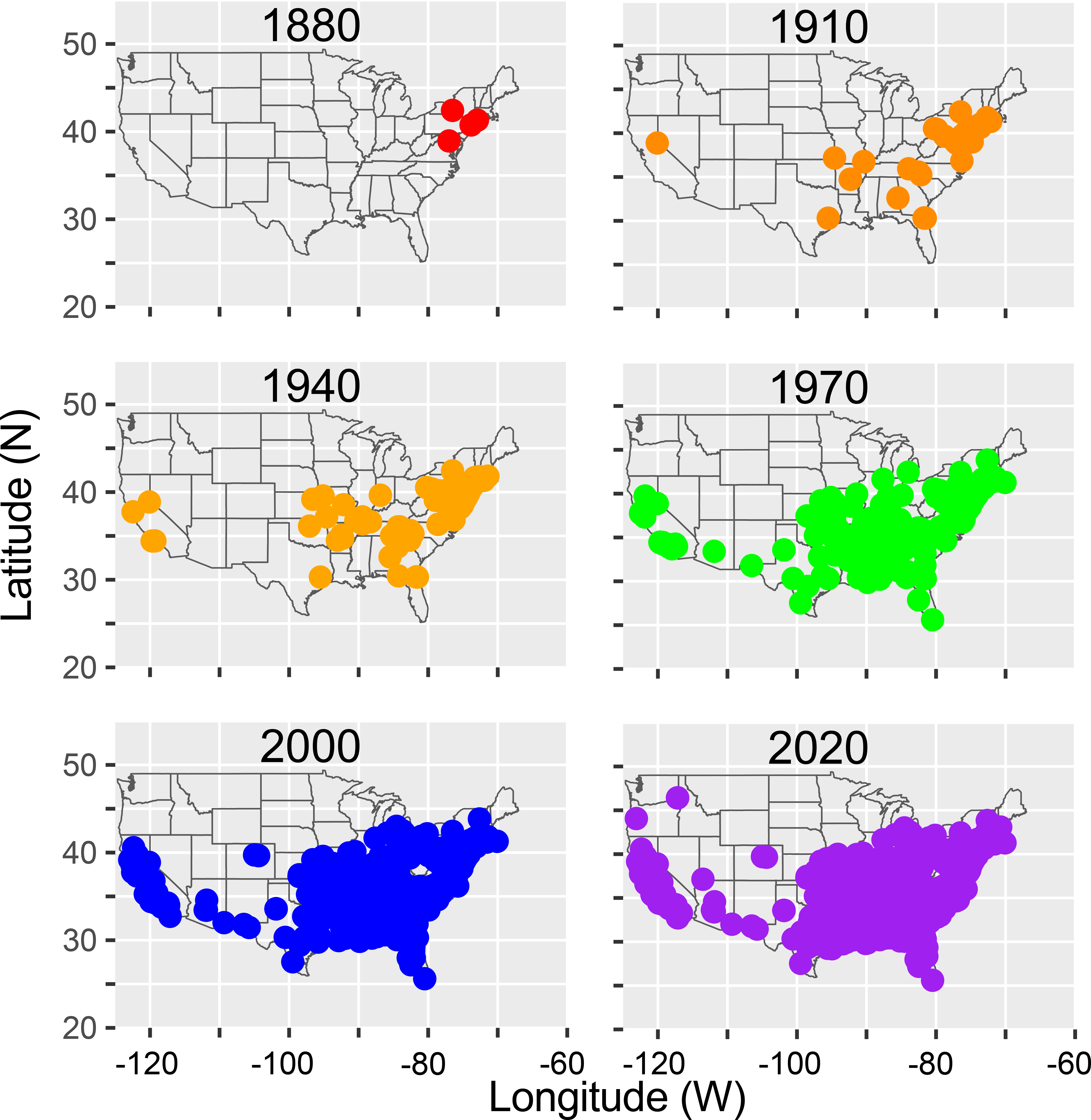
Maps of six time slices showing the invasion history of *Lonicera japonica* in the USA based on herbarium records. An animated version can be found at: https://github.com/barrettlab/2021-Genomics-bootcamp/wiki/2022-Biol-320-Lonicera-japonica-invasion-history-animation.

### Genetic diversity

After filtering and linkage disequilibrium thinning, 1,571 codominant SNP markers remained across 166 individuals, with 43.37% missing data (File S2, https://doi.org/10.5281/zenodo.7686073). Both observed and expected heterozygosity (*Ho* and *He*, respectively) were low across all sampling localities, with a mean *Ho* = 0.0319 and *He* = 0.1204. Mean inbreeding was relatively high (overall *Fis* = 0.7347). *He* ranged from 0.1-0.15, while *Ho* ranged from 0.2-0.45, and *Ho* was lower than *He* at all sampling localities (Fig. 2). *Ho* and *He* were highest at the Morgantown, Monongalia, WV and Preston, WV localities, and lowest at the Life Sciences Building and Deckers Creek sites (Monongalia, WV) localities; these four localities are all within a 20 km radius. Analysis of linkage disequilibrium after clone correction revealed significant values of *Ia* and *r-barD* at six localities (Table 1): Arboretum (Monongalia, WV), Jackson (WV), Lawrence (PA), Morgantown (Monongalia, WV), Preston (WV), and Star City (Monongalia, WV) (Table 1).

**Figure 2.**
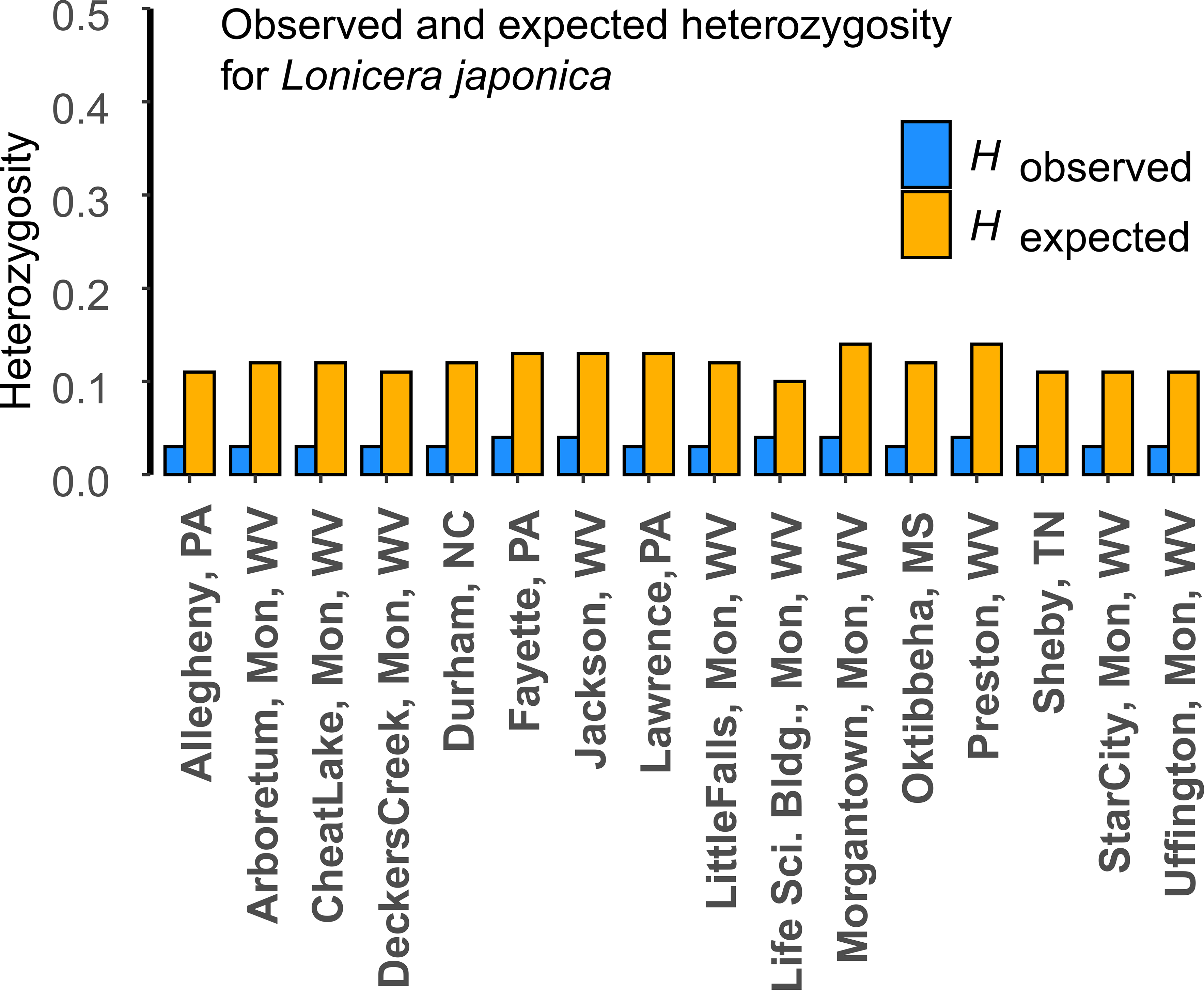
Estimates of observed (*Ho*, blue) and expected heterozygosity (*He*, orange) among sampling localities.

### Population structure

Analysis of molecular variance revealed a lack of overall population structure, with percentages of variation between localities = 0.38%, among individuals within localities = 0.31%, and within individuals = 99.3%, though none of the components of variation was significant. Principal Components Analysis of SNP data revealed an overall lack of clustering among sampling localities (Fig. 3A, B). Mean differentiation across localities was relatively low (overall *Fst* = 0.0347), and most pairwise comparisons of *Fst* between sampling localities ranged from 0-0.09, suggesting an overall lack of differentiation among populations (Fig. 3C). Discriminant Analysis of Principal Components (DAPC) revealed an optimal number of five genomic clusters (k = 5, optimal Principal Components retained = 80, Discriminant Functions retained = 3, BIC score = 624.98; Fig. 4A-C). Discriminant Axis 1 differentiated Cluster 5 from the remaining Clusters (Fig. 3A, B), and to a lesser extent differentiated Clusters 2 and 3. Discriminant Axis 2 further differentiated Cluster 2, 3, and 5, while Discriminant Axis 3 differentiated Clusters 1 and 4 (Fig. 3C). Plotting of ancestry coefficients from the DAPC revealed an overall pattern of population admixture, with all localities except for Fayette, Fig. 3D). Samples from the Arboretum locality PA composed of ≥2 genomic clusters ((Monongalia County, WV) were represented by all five clusters, whereas representatives of four clusters were observed in Morgantown (Monongalia, WV), Cheat Lake (Monongalia, WV), Shelby (TN), Little Falls (Monongalia, WV), Star City (Monongalia, WV), and Uffington (Monongalia, WV); it should be noted that samples sizes varied widely from each locality. Genomic Cluster 5 was only represented by a few individuals from the Arboretum locality, and was not sampled elsewhere in this study. Network analysis of multilocus genotypes reveal a similar overall pattern to the DAPC analysis, showing at least five genotype clusters, with no clear pattern of geographic structuring among them (Fig. 3E). Hierarchical clustering of Nei’s Genetic Distance among localities further supports an overall lack of population structure (Fig. 3F). Of all sampling localities, the Arboretum and Morgantown (Monongalia, WV) were most similar, followed by Preston (WV), Cheat Lake (Monongalia, WV), Decker’s Creek (Monongalia, WV), and Jackson (WV). A second cluster comprised three localities from Pennsylvania (Allegheny, Lawrence, and Fayette), Shelby (TN), and two localities from Monongalia, WV (Uffington and Life Sciences Building). The most divergent to all other localities was Oktibbeha (MS).

**Figure 3.**
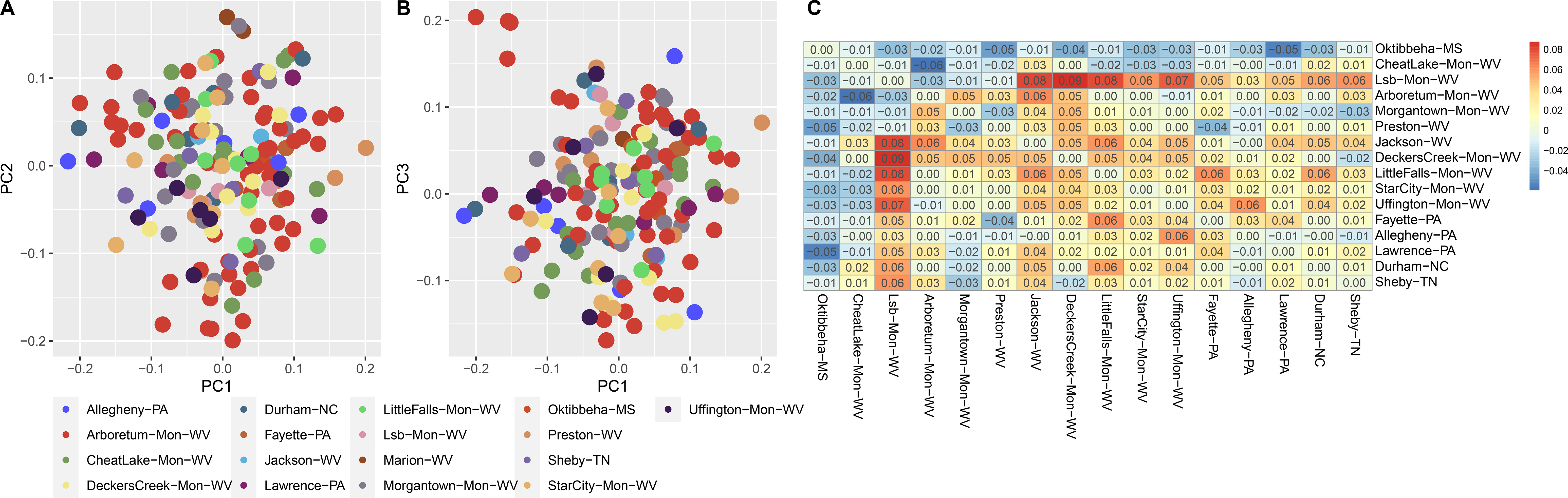
Population structure of *Lonicera japonica* across the eastern USA based on single nucleotide polymorphism (SNP) data. **A**. Principal Components Analysis (PCA) of 1,571 linkage disequilibrium-thinned SNPs, showing PCA Axes 1 and 2, and **B**. Axes 3 and 4. **C**. Pairwise *Fst* estimates among sampling localities.

**Figure 4.**
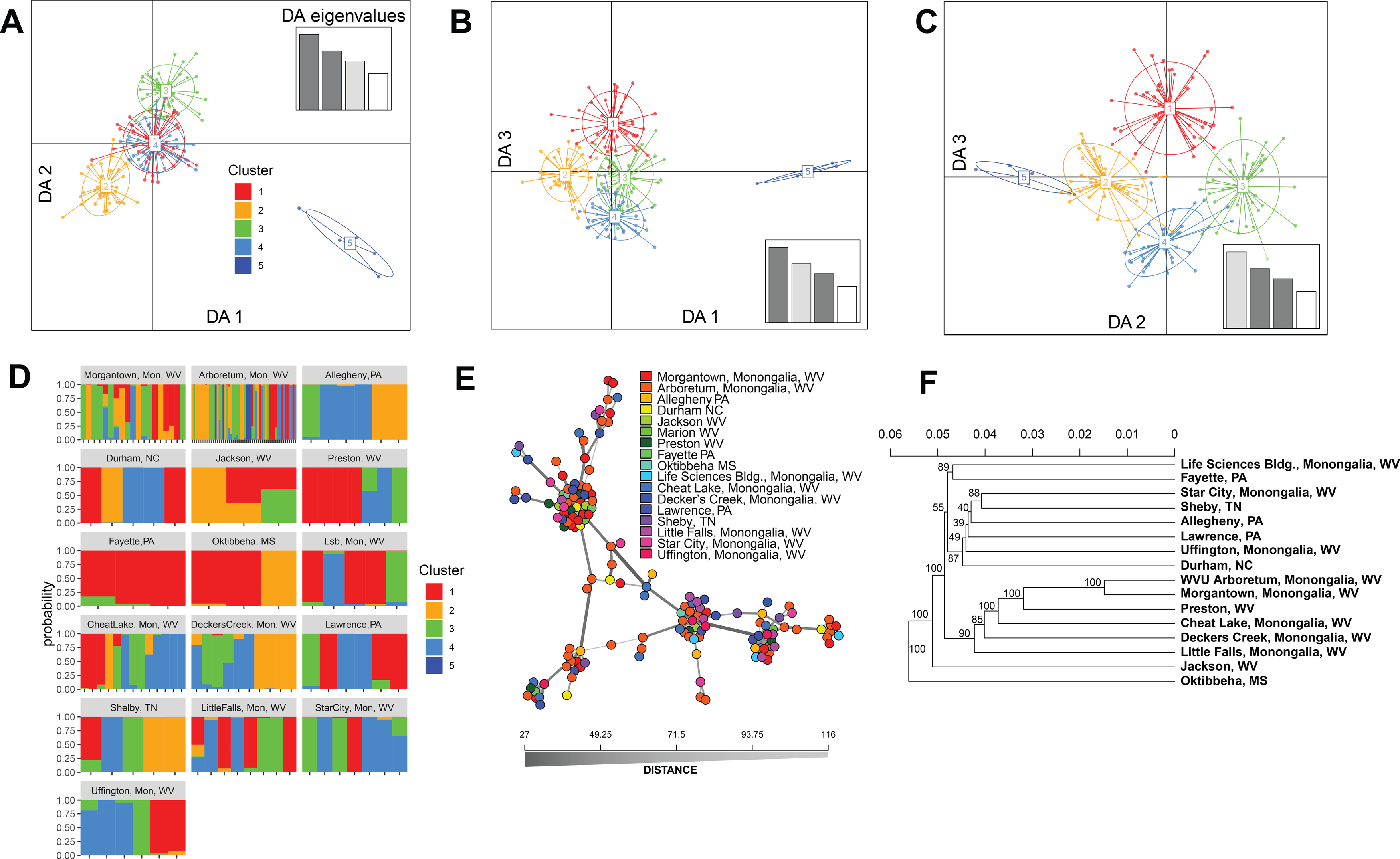
Additional analyses of population structure for *Lonicera japonica* in the eastern USA. **A-C**. Discriminant Analysis of Principal Components (DAPC) for the optimal value of k = 5. **A**. Axes 1 and 2, **B**. Axes 1 and 3, **C**. Axes 2 and 3. **D**. Ancestry coefficient plot of the five ancestral genomic population clusters by sampling locality. **E**. Multilocus genotype network, with colors in legend indicating sampling locality (origin) of each sample. **F**. Dendrogram based on Nei’s genetic distance depicting genetic distance among sampling localities.

## DISCUSSION

We conducted an analysis of invasion history, genetic variation, and population structure for one of the most problematic, weedy vines in eastern North America, *Lonicera japonica*. We reconstructed the invasion history of this species in the USA using digitized herbarium records, and showed the rapid spread across the USA in the early 1900s, including both regional spread and long distance dispersal to the western USA. We applied a cost-effective method for SNP genotyping (MIG-seq), leveraging a recently published chromosome-level genome sequence to quantify patterns of genomic variation. Our analyses revealed an overall lack of population structure and high inbreeding across the eastern USA, corroborating earlier studies based on allozymes, and in line with a species that was deliberately and ubiquitously introduced for erosion control and as an ornamental. Lastly, we discuss how we used plant invasion genomics as an accessible tool for integrating research and undergraduate education.

Invasion history of L. japonica in the USA

*Lonicera japonica* is thought to have been first introduced to the northeastern USA in 1862 (as the cultivar “Hall’s honeysuckle,” *L. japonica* var. *halliana*), but had likely arrived to the USA previously, as the earliest herbarium record of this species is from Kentucky in 1846 (Pelczar 1995; Schierenbeck 2004). Other records indicate that the species was present as far south as Virginia, Georgia, and Florida by 1900 (Leatherman 1955; Schierenbeck et al. 1995). Our analysis of digitized herbarium records corroborate these observations: by 1910, *L. japonica* was already widespread across the Northeast and Mid-Atlantic states, and had been collected in Georgia, Florida, the Carolinas, Missouri, Arkansas, and Texas, with a single record in California (Fig. 1). By 1970, it had become established in nearly all US states east of the Mississippi River, and was widespread throughout the eastern regions of Texas, Oklahoma, and Kansas. By 2000 it was widespread throughout nearly all of California at lower elevations, in southern Arizona and New Mexico, and in northeastern Colorado. Its continued expansion is evidenced by more recent collections in Oregon, Utah, and Idaho as of 2020.

Environmental constraints on the continued expansion of this species are believed to largely consist of soil characteristics (preferring well-drained, acidic soils), minimum winter temperatures (ice/frost damage), drought, and soil temperatures required for seed stratification (Leatherman 1955; Schierenbeck 2004). Interestingly, the earliest collections in the northeastern USA appeared close to the northern edge of the distribution in North America, although the species is now present in southern Maine, northern New York State, southern Ontario (Canada), northern Wisconsin, Michigan’s Upper Peninsula, and northwestern Washington state (https://www.eddmaps.org/species/subject.cfm?sub=3039, last accessed 24 February, 2023).

Continued northern and western expansion may be driven in part by milder winters and changing precipitation patterns due to climate change, but there is also evidence of adaptive evolution to withstand more extreme cold (Evans et al. 2013; Kilkenny and Galloway 2016). In common garden experiments, plants from the northern and western invasion fronts were less susceptible to cold than were plants from older, more established “core” regions of the invasive range in the US, suggesting post-establishment selection for cold tolerance along the invasion front (Kilkenny and Galloway 2016). In the eastern USA, *L. japonica* is predominantly found at lower elevations, but a similar adaptive scenario may be relevant to expansion in higher elevations (e.g. in Appalachia) as is the case for northward expansions (Strasbaugh and Core 1977; Hardt 1986; Pelczar 1995).

Genetic diversity and population structure of L. japonica in the USA

Analysis of 1,571 filtered SNPs and sampling of 166 individuals from 16 localities across the eastern USA for *L. japonica* revealed low overall genetic diversity and a general lack of population structure. Our overall estimate of population subdivision (*Fst* = 0.0347) was similar to that of Schierenbeck et al. (1995), who estimated *Gst* = 0.092 for localities sampled across eastern Georgia and western South Carolina based on allozyme data. Similarly, we found relatively high estimates of inbreeding coefficients (*Fis* overall = 0.7437), whereas Schierenbeck et al. calculated *Fis* =0.118 based on allozyme data. While these estimates are notably different, they both indicate some level of apparent inbreeding within populations, possibly driven in part by clonal propagation of this vining species, but also possibly driven by short dispersal distances for pollen (i.e. within patches). Possible explanations for these different *Fis* estimates may lie in differences in resolution between SNP and allozyme markers, our expanded sampling of populations across a broader geographic scale, temporal factors associated with ongoing neutral or adaptive evolution in this species (i.e. sampling in the 1990s vs 2020s), or some form of bias in either SNP-based or allozyme-based estimates. While we found no evidence for 100% identical clones at any of the sampling sites (Fig. 1E), we did find evidence of significant linkage disequilibrium at several sites (Table 1, based on significant values for *Ia* and *r-barD* after clone correction).

Our estimates of expected heterozygosity are also similar to Schierenbeck et al. (1995), with mean *He* = 0.1204 for SNP data vs. mean *He* = 0.189 for allozyme data, suggesting low overall levels of genetic diversity. Other representations of population structure further demonstrate a lack of distinctness among localities across the broader eastern-US invasive range for this species (PCA, DAPC, multilocus genotype network analysis, hierarchical cluster analysis; Fig. 4). There is no discernable pattern of geographic differentiation evident from our analyses, suggesting a highly admixed gene pool for this species in North America. It must be noted that our main sampling focus was in northern West Virginia (mid-latitude), and thus our interpretation of overall patterns of variation may have been influenced by this. However, this does not appear to be the case, as sampling localities in WV are virtually indistinguishable from those more broadly sampled in the eastern US (Fig. 4 D-F).

Many factors may have contributed to the patterns observed in this and earlier studies of genetic variation in *L. japonica*. First, this is an obligately outcrossing, perennial species, pollinated by a variety of animals including birds and both diurnal and nocturnal insects (Leatherman 1955; Miyake and Yahara 1998). Further, seeds are dispersed locally or perhaps more broadly by mammals and birds (e.g. White and Styles 1982). Both the pollination and seed dispersal syndromes of this species would be expected to lead to frequent local or regional dispersal, effectively facilitating admixture (e.g. Bariball et al. 2015). More importantly, however, is the way in which *L. japonica* was likely spread across the USA, both historically and contemporarily. Repeated anthropogenic dispersal by deliberate planting for erosion control and ornamental purposes, followed by local escapes from cultivation and subsequent spread may be a stronger factor in determining the current distribution of genetic variation in this species than wildlife-mediated dispersal (e.g. Brusa and Holzapfel 2018; Alvarado-Serrano et al. 2019). Thus, it is not surprising that *L. japonica* would exhibit low levels of population structure in the eastern US, especially given its long history as an invasive species in the USA and the fact that it is still sold in garden stores (Schierenbeck 2004; C. Barrett *personal observation*). By comparison, the few studies on population genetics of *L. japonica* in the native range suggest relatively higher levels of population structure (e.g. Fu et al. 2013; He et al. 2016; He et al. 2017), but these studies likely included multiple, possibly divergent varieties that may not be represented in the USA. Certainly, future research on population genomics of *L. japonica* should seek to sample representatives across the entire spectrum of variation in the native and invasive ranges (the former including representatives of all known varieties, and the latter on multiple continents), to compare patterns of genetic variability and trace the origins of invasive populations.

Traditional theory of invasion genetics centered around the expectation of single introductions, drastic genetic bottlenecks upon establishment representing a fraction of the diversity from the native range, and subsequent spread (e.g. see Barrett and Husband 1990; Novak 2007; Dlugosch and Parker 2008; Barrett 2015). Yet, as genomic methods enable a rapid increase in invasion studies, the patterns emerging are not so simple, and the aforementioned scenario seems to be the exception rather than the rule (e.g. Sakai et al. 2001; Lee 2002; Frankham 2005; Dlugosch and Parker 2008; Sutherland et al. 2021). For example, several studies have concluded that invasions are often repeated events, with multiple introductions, subsequent establishments, spread from points of introduction, secondary contact, and possibly hybridization with native or other invasive species (Ellstrand and Schierenbeck 2000, Blair and Hufbauer 2010). *Lonicera japonica* would be expected to fall somewhere in the latter category, given the overall pattern of admixture and lack of population structure observed, which was likely facilitated and exacerbated by deliberate, human-aided dispersal over two centuries (Figs. 3, 4). Admixture after multiple invasions may provide genetic variation for rapid adaptation to conditions in the invasive range, bringing together novel allelic combinations that otherwise would have remained geographically isolated in the native range (Dlugosch et al. 2015). *Lonicera japonica* represents an apt case study in global patterns of rapid evolution, as it is invasive on all continents aside from Antarctica (Schierenbeck 2004). Spatiotemporal comparisons of patterns of invasion history and genomic variation (e.g. using material sampled from herbarium specimens) will be extremely powerful in elucidating the environmental and genomic factors associated with rapid, post-invasion evolution (e.g. Kreiner et al. 2022). Future studies should emphasize collecting densely sampled SNP data from populations in the native and invasive ranges (on a global scale) to identify: 1) fundamental differences in population structure and genetic diversity in the native vs. invasive ranges; 2) spatiotemporal patterns of variation linked to invasion routes and invasion history; and 3) evidence for adaptive variation linked to climate, soils, pathogens (or a lack thereof), and other environmental factors post-invasion.

### Using plant invasion genomics to integrate teaching and research

This study is the product of a senior capstone course in the Department of Biology at West Virginia University, *Biology 320 Total Science Experience: Genomics*. The course’s major objective was to allow students to experience the entire scientific process from start to finish, with the following objectives: 1) identify a research question, 2) develop a testable hypothesis, 3) write a proposal to fund the research, 4) conduct the research itself, 5) write a scientific paper on the findings and interpretation their meaning, and 6) present a poster at an end-of-semester symposium. We chose to focus our hypotheses on a single species, *Lonicera japonica*, which in WV retains at least some leaves throughout the winter (the Spring Semester starts in mid-January so students could collect material early on), is highly abundant and invasive locally, is relatively easy for students to identify, and requires no collecting permits or Institutional Board Review (e.g. for working with animals).

Our choice to focus on a single species (*L. japonica*) represented a “happy medium” between allowing all student groups to choose their own organisms/hypotheses of interest freely, and dictating projects directly to the students, neither of which we felt would maximize the learning experience. The added bonus of our approach was to generate a product as a class that could be used for specific comparisons to the data and hypotheses tested within each group, and further be submitted for publication with all students as coauthors (Appendix 1). Lab and bioinformatic protocols (conducted primarily in the R computing environment) were made freely available to the students and anyone else interested in gaining research experience or taking a similar approach in a lab-based course via the GitHub wiki. This provided a “one-stop shop” of sorts for data access, R commands, and tutorials that users anywhere can follow. For example,: https://github.com/barrettlab/2021-Genomics-bootcamp/wiki/2022-Biol-320-Migseq-data-processing-commands-on-myco-server, https://github.com/barrettlab/2021-Genomics-bootcamp/wiki/2022-Biol-320-Population-genomic-analyses-with-R.

The goal was to allow student groups to develop their own “semi-directed” hypotheses (i.e. within the contexts of invasion genomics and landscape genetics), plan their sampling around those hypotheses, and in the end contribute their data to a class-wide analysis. For example, one group’s hypothesis was: “We expect *Lonicera japonica* to have higher genetic diversity in more disturbed vs. less disturbed habitats.” Students worked in groups of three, and were each responsible for collecting leaf material of *L. japonica*, supplemented by collections from colleagues across the central and eastern USA (listed in the Acknowledgements section). Students learned: 1) identification of common invasive plant species in the eastern USA (https://www.youtube.com/watch?v=zOfiXoy9BIA); 2) background information on genomics, bioinformatics, population genetic theory/practice, and invasion biology; 3) experimental design; 4) wet-lab skills; 5) population-genetic data analysis in the R programming language; 6) how to conduct literature searches; 7) effective proposal and manuscript writing; and 8) effective oral communication skills.

As final capstone products, students: 1) produced group-written research proposals following general proposal guidelines (i.e. the US National Science Foundation), with frequent undergraduate (peer), graduate, and faculty reviews of their work; 2) presented their findings for the general public and students in other capstone courses in an undergraduate research symposium; and 3) produced manuscripts that challenged them to write beyond standard lab reports. Based on end-of-semester student feedback, their experiences were generally positive with most having gained a deeper appreciation and understanding of the research process as a whole. While we did not formally test students’ overall understanding of experimental design, e.g. The Experimental Design Ability Test (EDAT; Sirum and Humburg 2011), we observed significant improvements in students’ communication, wet-lab, and data-analysis skills based on weekly formative assessments throughout the semester (e.g. proposal feedback, peer reviews, homework activities, and manuscript feedback). Certainly EDAT or a similar summative assessment should be implemented in future offerings of the course more formally and explicitly.

In summary, plant invasion genomics was used as an engaging, accessible, and novel way to provide meaningful research experience for undergraduate biology students.

## ACKNOWLEDGEMENTS

Research funding was provided by the West Virginia University Biology Department, and by US National Science Foundation Awards OIA-1920858 to C. Barrett and DEB-1542509 to S. DiFazio. We thank the following colleagues for collecting material across the USA: Bonnie Isaac, Mason Heberling, Jennifer Mandel, James Beck, Vanessa Koelling, Claudia Stein, Mark Fishbein, Paul Manos, and Ryan Folk. We thank Michael McKinstry and Ryan Percifield (WVU Genomics Core Facility) for lab and sequencing assistance. We acknowledge support from the WVU Genomics Core Facility, and CTSI Grant #U54 GM104942 which provides financial support to the Core Facility.

## APPENDICES

### Appendix 1, Spring 2022 Biology 320 Students

Ashkanani, Ali; Bauman, Gabrielle; Brooks, David; Brooks, Zane; Butler, Zye; Carder, Cayton; Carter, Marissa; Collins, Samuel; Cumberledge, Aubrey; Daniels, Kellie; Deery, Rachel; Duncan, Taylor; England, Payton; Fergione, Alexander; French, Daze; Fuchs, Taylor; Garris, Megan; Harris, Alexander; Hughes, Christopher; Katz, Morgan; Komatsu, Ryusa; Lalama, Madelyn; Limer, Kaley; Litten, Zoe; Long, Ryan; Lowther, Alanna; Marku, Vjosa; McGraw, Aidan; Midkiff, Rachel; Mitchell, Victor; Parascandola, Lowell; Pool, Hunter; Riel, Stephanie; Runion, Anna; Safar, Ammar; Setser, Amelia; Shifflett, Victoria; Smith, Jamie; Smothers, Jacob; Stover, Cassandra; Tarsi, Tavia; Taylor, Mark; Tooker, Tiffany; Walls, Austin; White- Montgomery, Katielee; Wunder, Michelle; Wyllie, Heather.

## Notes

https://doi.org/10.5281/zenodo.7686073

https://github.com/barrettlab/2021-Genomics-bootcamp/wiki/2022-Biol-320-Lonicera-japonica-invasion-history-animation

https://github.com/barrettlab/2021-Genomics-bootcamp/wiki/2022-Biol-320-Migseq-data-processing-commands-on-myco-server

https://github.com/barrettlab/2021-Genomics-bootcamp/wiki/2022-Biol-320-Population-genomic-analyses-with-R

### Competing Interest Statement

The authors have declared no competing interest.

