## Supplementary material for "A lack of population structure characterizes the invasive *Lonicera japonica* in West Virginia and across eastern North America": File S1

**PCR1 – multiplex – modified 16s amplicon primers**

Forward primers: i5 Nextera Pad + N or NN spacer + SSR + **anchor**

(ACT)$_4$TG-f-16S-1(2)N TCGTCGGCAGCGTCAGATGTGTATAAGAGACAG N(NN)ACTACTACTACTTG
(CTA)$_4$TG-f-16S-1(2)N TCGTCGGCAGCGTCAGATGTGTATAAGAGACAG N(NN)CTACTACTACTATG
(TTG)$_4$AC-f-16S-1(2)N TCGTCGGCAGCGTCAGATGTGTATAAGAGACAG N(NN)TTGTTGTTGTTGAC
(GTT)$_4$CC-f-16S-1(2)N TCGTCGGCAGCGTCAGATGTGTATAAGAGACAG N(NN)GTTGTTGTTGTTCC
(GTT)$_4$TC-f-16S-1(2)N TCGTCGGCAGCGTCAGATGTGTATAAGAGACAG N(NN)GTTGTTGTTGTTTC
(GTG)$_4$AC-f-16S-1(2)N TCGTCGGCAGCGTCAGATGTGTATAAGAGACAG N(NN)GTGGTGGTGGTGAC
(GT)$_6$TC-f-16S-1(2)N TCGTCGGCAGCGTCAGATGTGTATAAGAGACAG N(NN)GTGTGTGTGTGTTC
(TG)$_6$AC-f-16S-1(2)N TCGTCGGCAGCGTCAGATGTGTATAAGAGACAG N(NN)TGTGTGTGTGTGAC

Reverse primers: i7 Nextera Pad + N or NN spacer + SSR + **anchor**

(ACT)$_4$TG-r-16S-1(2)N GTCTCGTGGGCTCGGAGATGTGTATAAGAGACAG N(NN)ACTACTACTACTTG
(CTA)$_4$TG-r-16S-1(2)N GTCTCGTGGGCTCGGAGATGTGTATAAGAGACAG N(NN)CTACTACTACTATG
(TTG)$_4$AC-r-16S-1(2)N GTCTCGTGGGCTCGGAGATGTGTATAAGAGACAG N(NN)TTGTTGTTGTTGAC
(GTT)$_4$CC-r-16S-1(2)N GTCTCGTGGGCTCGGAGATGTGTATAAGAGACAG N(NN)GTTGTTGTTGTTCC
(GTT)$_4$TC-r-16S-1(2)N GTCTCGTGGGCTCGGAGATGTGTATAAGAGACAG N(NN)GTTGTTGTTGTTTC
(GTG)$_4$AC-r-16S-1(2)N GTCTCGTGGGCTCGGAGATGTGTATAAGAGACAG N(NN)GTGGTGGTGGTGAC
(GT)$_6$TC-r-16S-1(2)N GTCTCGTGGGCTCGGAGATGTGTATAAGAGACAG N(NN)GTGTGTGTGTGTTC
(TG)$_6$AC-r-16S-1(2)N GTCTCGTGGGCTCGGAGATGTGTATAAGAGACAG N(NN)TGTGTGTGTGTGAC

**PCR2 – i5 and i7 Nextera sequences: Adapter + barcode + 16S Nextera pad**

**I5-Locus Primer binding for 2 step** **TCGTCGGCAGCGTC**
**Example 16S-F** **TCGTCGGCAGCGTCAGATGTGTATAAGAGACAGNNNNCCTACGGGNGGCWGCAG**

i5 (forward) oligos

I5Amp-01 AATGATACGGCGACCACCGAGATCTACACAAACGGTCGTCGGCAGCGTC
I5Amp-02 AATGATACGGCGACCACCGAGATCTACACACATACTCGTCGGCAGCGTC
I5Amp-03 AATGATACGGCGACCACCGAGATCTACACACTCTTTCGTCGGCAGCGTC

| | |
|---|---|
| I5Amp-04 | AATGATACGGCGACCACCGAGATCTACACATGTTGTCGTCGGCAGCGTC |
| I5Amp-05 | AATGATACGGCGACCACCGAGATCTACACATTCCGTCGTCGGCAGCGTC |
| I5Amp-06 | AATGATACGGCGACCACCGAGATCTACACCCTAACTCGTCGGCAGCGTC |
| I5Amp-07 | AATGATACGGCGACCACCGAGATCTACACCGAGGCTCGTCGGCAGCGTC |
| I5Amp-08 | AATGATACGGCGACCACCGAGATCTACACCTTTGCTCGTCGGCAGCGTC |
| I5Amp-09 | AATGATACGGCGACCACCGAGATCTACACGAAATGTCGTCGGCAGCGTC |
| I5Amp-10 | AATGATACGGCGACCACCGAGATCTACACGCTCAATCGTCGGCAGCGTC |
| I5Amp-11 | AATGATACGGCGACCACCGAGATCTACACGGACTTTCGTCGGCAGCGTC |
| I5Amp-12 | AATGATACGGCGACCACCGAGATCTACACGTTGGATCGTCGGCAGCGTC |
| I5Amp-13 | AATGATACGGCGACCACCGAGATCTACACTAAGCTTCGTCGGCAGCGTC |
| I5Amp-14 | AATGATACGGCGACCACCGAGATCTACACTCTCGGTCGTCGGCAGCGTC |
| I5Amp-15 | AATGATACGGCGACCACCGAGATCTACACTCTTCTTCGTCGGCAGCGTC |
| I5Amp-16 | AATGATACGGCGACCACCGAGATCTACACTTTGTCTCGTCGGCAGCGTC |

**I7-Locus Primer binding for 2 step** GTCTCGTGGGCTCGG

**Example 16S-R** GTCTCGTGGGCTCGGAGATGTGTATAAGAGACAGNNNNGACTACHVGGGTATCTAATCC

i7 (reverse) oligos

| | |
|---|---|
| I7Amp-01 | CAAGCAGAAGACGGCATACGAGATCGTGATGTCTCGTGGGCTCGG |
| I7Amp-02 | CAAGCAGAAGACGGCATACGAGATACATCGGTCTCGTGGGCTCGG |
| I7Amp-03 | CAAGCAGAAGACGGCATACGAGATGCCTAAGTCTCGTGGGCTCGG |
| I7Amp-04 | CAAGCAGAAGACGGCATACGAGATTGGTCAGTCTCGTGGGCTCGG |
| I7Amp-05 | CAAGCAGAAGACGGCATACGAGATCACTGTGTCTCGTGGGCTCGG |
| I7Amp-06 | CAAGCAGAAGACGGCATACGAGATATTGGCGTCTCGTGGGCTCGG |
| I7Amp-07 | CAAGCAGAAGACGGCATACGAGATGATCTGGTCTCGTGGGCTCGG |
| I7Amp-08 | CAAGCAGAAGACGGCATACGAGATTCAAGTGTCTCGTGGGCTCGG |
| I7Amp-09 | CAAGCAGAAGACGGCATACGAGATCTGATCGTCTCGTGGGCTCGG |
| I7Amp-10 | CAAGCAGAAGACGGCATACGAGATAAGCTAGTCTCGTGGGCTCGG |
| I7Amp-11 | CAAGCAGAAGACGGCATACGAGATGTAGCCGTCTCGTGGGCTCGG |
| I7Amp-12 | CAAGCAGAAGACGGCATACGAGATTACAAGGTCTCGTGGGCTCGG |
| I7Amp-13 | CAAGCAGAAGACGGCATACGAGATTTGACTGTCTCGTGGGCTCGG |
| I7Amp-14 | CAAGCAGAAGACGGCATACGAGATGGAACTGTCTCGTGGGCTCGG |
| I7Amp-15 | CAAGCAGAAGACGGCATACGAGATTGACATGTCTCGTGGGCTCGG |
| I7Amp-16 | CAAGCAGAAGACGGCATACGAGATACGTTGGTCTCGTGGGCTCGG |
| I7Amp-17 | CAAGCAGAAGACGGCATACGAGATCGTAGCGTCTCGTGGGCTCGG |
| I7Amp-18 | CAAGCAGAAGACGGCATACGAGATGTACCAGTCTCGTGGGCTCGG |
| I7Amp-19 | CAAGCAGAAGACGGCATACGAGATTACGATGTCTCGTGGGCTCGG |
| I7Amp-20 | CAAGCAGAAGACGGCATACGAGATGCAGGCGTCTCGTGGGCTCGG |
| I7Amp-21 | CAAGCAGAAGACGGCATACGAGATCATAAGGTCTCGTGGGCTCGG |
| I7Amp-22 | CAAGCAGAAGACGGCATACGAGATATGTTAGTCTCGTGGGCTCGG |
| I7Amp-23 | CAAGCAGAAGACGGCATACGAGATTGCGGTGTCTCGTGGGCTCGG |
| I7Amp-24 | CAAGCAGAAGACGGCATACGAGATGAGGTTGTCTCGTGGGCTCGG |

PCR1 + PCR2 Example:

$(ACT)_4$TG-f-16S-1(2)N TCGTCGGCAGCGTCAGATGTGTATAAGAGACAG N(NN)ACTACTACTACTTG

I5Amp-01 AATGATACGGCGACCACCGAGATCTACACAAACGGTCGTCGGCAGCGTC

(ACT)$_4$TG-r-16S-1(2)N GTCTCGTGGGCTCGGAGATGTGTATAAGAGACAG N(NN)ACTACTACTACTTG

I7Amp-01 CAAGCAGAAGACGGCATACGAGATCGTGATGTCTCGTGGGCTCGG

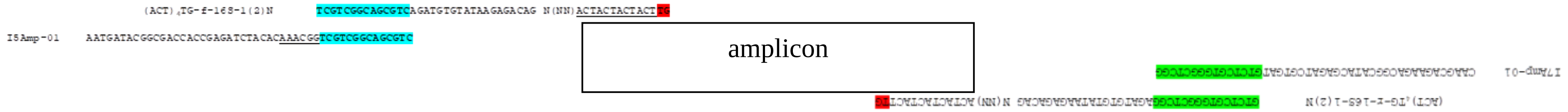
